# Deconvolving multiplexed histone modifications in single cells

**DOI:** 10.1101/2021.04.26.440629

**Authors:** Jake Yeung, Maria Florescu, Peter Zeller, Buys Anton de Barbanson, Alexander van Oudenaarden

## Abstract

Recent advances have enabled mapping of histone modifications in single cells^1–12^, but current methods are constrained to profile only one histone modification per cell. Here we present an integrated experimental and computational framework, scChIX (single-cell chromatin immunocleavage and unmixing), to map multiple histone modifications in single cells. scChIX multiplexes two histone modification profiles together in single cells, then computationally deconvolves the signal using training data from respective histone modification profiles. We first validate this method using purified blood cells and show that although the two repressive marks, H3K27me3 and H3K9me3, are generally mutually exclusive, the transitions between the two regions can vary between cell types. Next we apply scChIX to a heterogenous cell population from mouse bone marrow to generate linked maps of active (H3K4me1) and repressive (H3K27me3) chromatin landscapes in single cells, where coordinates in the active modification map correspond to coordinates in the repressive map. Linked analysis reveals that immunoglobulin genes in the *Igkv* region are in a repressed chromatin state in pro-B cells, but become activated in B cells. Overall, scChIX unlocks systematic interrogation of the interplay between histone modifications in single cells.

## Introduction

Large-scale efforts characterizing different histone modifications in a variety of cell populations commonly use chromatin immunoprecipitation followed by sequencing (ChIP-seq)^10, 11, 13–16^. Alternative strategies to ChIP-seq based on enzyme tethering (chromatin immunocleavage, ChIC) have reduced the background signal in profiling the epigenome^17^, and have enabled single-cell profiling of histone modifications^1–9, 12^. Tethering strategies involve incubating cells with an antibody against a histone modification of interest, which then tethers either protein A-MNase^1, 3, 9, 12^ or protein A-Tn5^2, 4–8^ fusion protein to generate targeted fragments in single cells. However, techniques to map single-cell histone modifications are currently limited to only one histone modification per single cell.

We present an integrated experimental and computational framework for multiplexing histone modifications in single cells. To profile two histone modifications in single cells (Figure 1a), we first generate three genome-wide sortChIC^9^ datasets: two datasets by incubating cells with one of the two histone modification antibodies separately (single-incubated, Figure 1b), and the third by incubating cells with both histone modification antibodies together (double-incubated, Figure 1b). We then use our two single-incubated datasets as training data to generate the possible pairs of genome-wide histone modification profiles that, when added together, fit to a single-cell profile from the double-incubated dataset (Figure 1c). For each double-incubated cell, we then deconvolve the multiplexed data by probabilistically assigning each fragment back to their respective histone modification.

**Figure 1.**
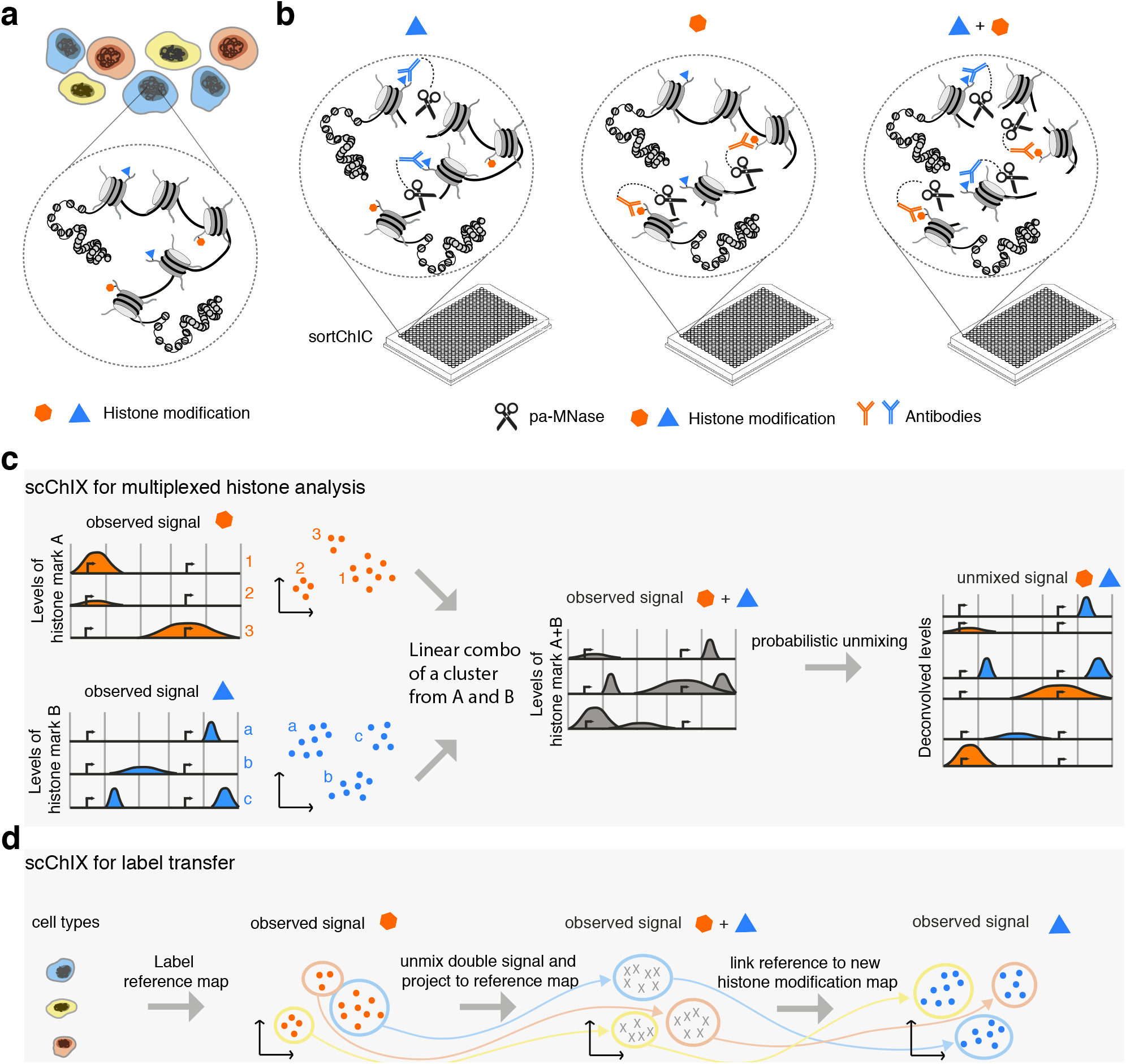
Overview of the scChIX method. **a** Chromatin regulation of different cell types (different colored cells) is regulated in part through multiple histone modifications (two histone modifications shown as an example). **b** Input for scChIX are three sortChIC antibody incubation conditions: two conditions each target a single histone modification (single-incubated) only and the third condition targeting both histone modifications simultaneously (double-incubated). **c** Schematic of scChIX for deconvolving multiplexed histone modifications. The two single-incubated sortChIC datasets (one targeting histone modification A, the other modification B) are training data to define the possible pairs of histone modification distributions that can be combined to generate a hypothetical double-incubated cell. For each observed double-incubated cell, we then assign the cell to the most likely pair of cell states, one from each histone modification. We then probabilistically assign each pA-MNase cut into their respective histone modification. **d** Label transfer allows joint analysis of two single-incubated sortChIC datasets targeting functionally distinct histone modifications. Information derived from one histone modification, such as cell types, histone mark levels, and pseudotime, can be transferred to another histone modification, using the double-incubated cells as a link.

scChIX also links chromatin regulation maps of distinct histone modifications, integrating different chromatin states in single cells. Locations in the single-cell map for one histone modification directly correspond to locations for another histone modification. In these linked maps, information derived from one chromatin state, such as cell types, histone mark levels, and pseudotimes, can transfer to another chromatin state (Figure 1d), unlocking joint analysis of multiple histone modifications in single cells.

## Results

### Ground truth data validates scChIX method

To validate our method, we generate a ground truth sortChIC dataset by purifying three known cell types from mouse bone marrow: B cells, granulocytes, and NK cells, using FACS and applying scChIX (Methods). Of note, the sortChIC method is designed to integrate FACS with histone modification mapping^9^, so we can enrich for a cell type and map histone modifications in one workflow. We split bone marrow cells into three technical batches. We incubate one batch with anti-H3K27me3 antibody alone (single-incubated), one with anti-H3K9me3 alone (single-incubated), and finally one batch with both anti-H3K27me3 and anti-H3K9me3 antibodies together (double-incubated, H3K27me3+H3K9me3). We then sort cells into 384-well plates, each plate containing all three cell types, and generate targeted cut fragments.

*A priori*, we do not know, from the double-incubated data alone, which cut fragments correspond to H3K27me3 and which to H3K9me3. We observe only a superposition of the two profiles. We therefore use the single-incubated sortChIC data to train a statistical model of how cells from the same cell type combine their H3K27me3 and H3K9me3 profiles to generate double-incubated cut fragments. This model is then used to deconvolve the single-cell multiplexed signal into their respective histone modifications (Methods).

We apply latent dirichlet allocation (LDA)^18^, which factorizes count matrices based on a multinomial model (Methods), to each of the three sortChIC datasets (Supplemental Figure 2a). The sortChIC datasets show separation between the FACS-identified cell types, confirming cell type-specific distributions of repressive histone modifications. Although we find two subclusters of NK cells, which could arise from heterogeneity from FACS enrichment, we could still use the NK cell identities derived from FACS to validate scChIX. Downstream analysis of LDA finds clusters defined by the topic weights (Methods, Supplemental Figures 2b,c). We use these topic weights to cluster cells in the H3K27me3 and H3K9me3 sortChIC datasets (Supplemental Figure 2a, left two panels).

**Figure 2.**
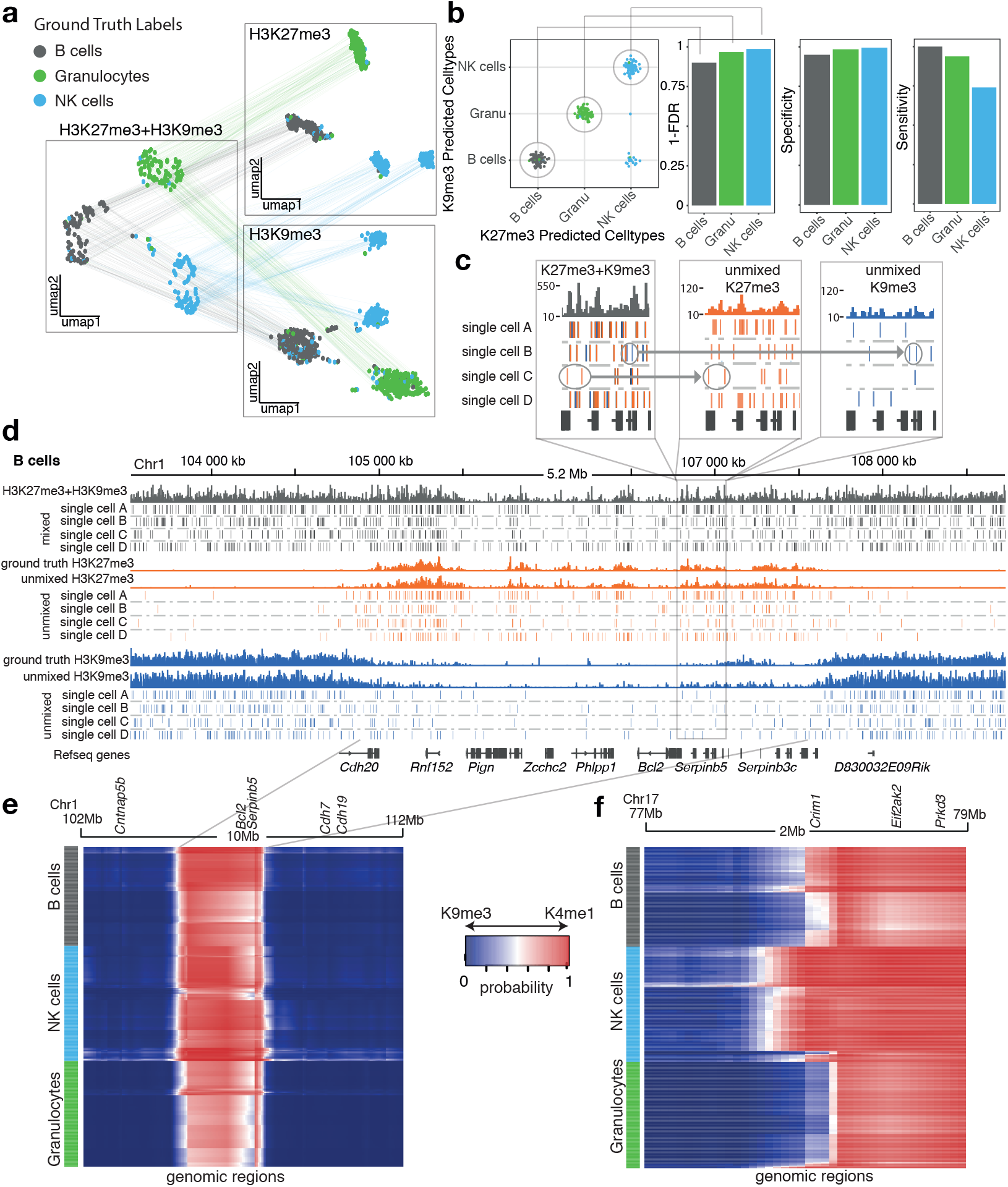
scChIX accurately deconvolves multiplexed histone modifications in single cells. **a** UMAP representation of the H3K27me3 and H3K9me3 histone modification space derived from the two single-incubated datasets (right two panels), and the H3K27me3+H3K9me3 space (left panel) derived from the double-incubated data. Cells are colored by their ground truth cell type labels. The cells in the H3K27me3- and H3K9me3-only space have unmixed double-incubated cells whose deconvolved signal has been projected onto their respective UMAPs. Lines connecting across datasets connect where each double-incubated cell is located in each of the three histone modification space. **b** Matrix summarizing the cluster pair that scChIX selected for each double-incubated cell. Cells along the diagonal are predicted to be B cells, Granulocytes, and NK cells, respectively. Cells in the off-diagonal are false negatives. Barplots summarizing false discovery rate (FDR), sensitivity, and specificity of assigning each cell type (right). **c** Zoom-in coverage plot and single-cell cut fragments in B cells of mixed (H3K27me3+H3K9me3, grey bars), unmixed (H3K27me3 and H3K9me3, orange and blue bars). Positions of cut fragments are shown for four single cells (single cells A, B, C, and D) for H3K27me3+H3K9me3 signal (grey ticks) as well as their unmixed outputs (orange and blue ticks). Circled reads and arrow highlight examples of cut fragments being assigned to either H3K27me3 (orange) or H3K9me3 (blue). **d** Zoom-out of the *Serpinb5* locus. Cut fragments from HK27me3+K9me3 are colored based on whether they have been assigned to H3K27me3 (orange) or H3K9me3 (blue). Ground truth coverage are single-incubated sortChIC data targeting H3K27me3 (orange) and H3K9me3 (blue). **e** Heatmap of probabilities *p* of assigning reads to H3K27me3 (*p* = 1, red) or H3K9me3 (*p* = 0, blue) around the *Bcl2* locus. Rows are single cells (ordered by predicted cell type), columns are genomic regions (50 kilobase bins). Transitions between H3K9me3- and H3K27me3-marked chromatin states are independent of cell type. **f** Same as (e) but at the *Crim1* locus, where transitions from H3K9me3 to H3K27me3 (blue to red) are cell type-specific.

Demultiplexing the double-incubated data involves two steps (Methods). First, we use the training data to infer which genome-wide H3K27me3 distribution was added to which H3K9me3 distribution to generate a mixture of two distributions (H3K27me3+H3K9me3). Second, we probabilistically assign each double-incubated cut fragment to either H3K27me3 or H3K9me3, given that we know the underlying mixture of the two profiles.

The deconvolved H3K27me3+H3K9me3 data generates two sets of cuts for each cell: one set coming from H3K27me3 and the other from H3K9me3. We project the two sets of cuts onto the H3K27me3 or H3K9me3 latent space (derived from LDA), respectively (Figure 2a). Since each deconvolved cell has a set of cuts in H3K27me3 and H3K9me3 simultaneously, we can link the UMAPs together, creating a joint chromatin regulation space (Figure 2a).

The double- and single-incubated cells in the H3K27me3 and H3K9me3 UMAPs intermingle, suggesting that the model accurately assigns cut fragments to their respective histone modification (Supplemental Figure 3a, b). Comparing the H3K27me3 deconvolved pseudobulk signal with our ground truth single-incubated pseudobulk shows high correlation for the expected cell type, and lower for the other two cell types (Supplemental Figure 3c). The H3K9me3 deconvolved pseudobulk signal also shows highest correlation with the expected cell type, with lower correlation from other cell types (Supplemental Figure 3d). Overall, our ground truth dataset demonstrates that scChIX is accurate in assigning pA-MNase cuts to their respective histone modification.

**Figure 3.**
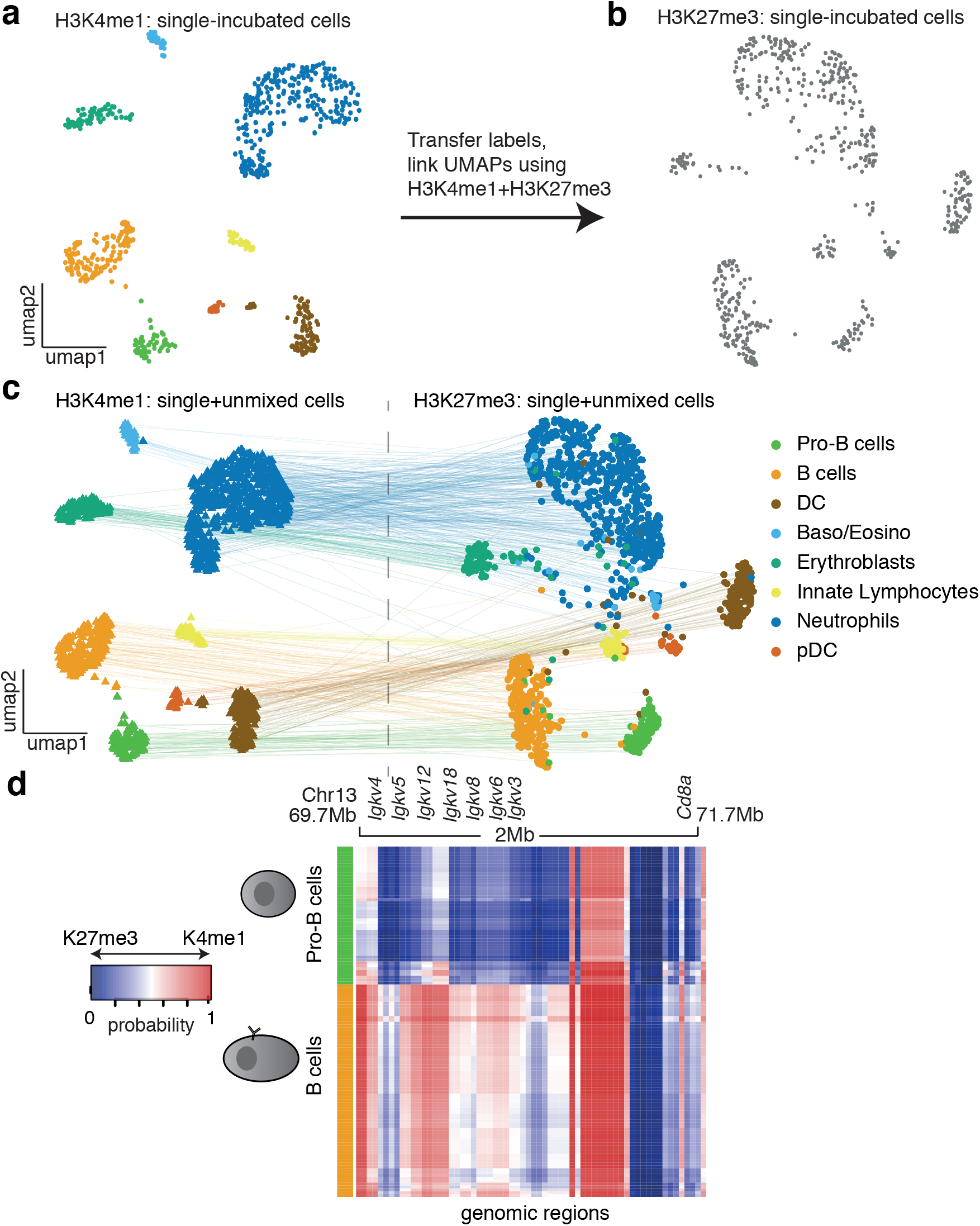
scChIX enables joint analysis of distinct histone modifications in single cells. **a** UMAP of sortChIC signal of H3K4me1 in bone marrow. Clusters are colored by cell type. **b** UMAP of sortChIC signal of H3K27me3 in whole bone marrow. Cell types in H3K27me3 are inferred by transferring labels from H3K4me1. **c** H3K4me1 and H3K27me3 UMAPs linked together by deconvolved double-incubated cells. H3K4me1 and H3K27me3 portions of the double-incubated cells are projected onto their respective UMAPs. Lines connect where the active signal and the corresponding repressive signal are located for each double-incubated cell. **d** Heatmap of probability of assigning a read in a region to either H3K27me3 or H3K4me1. Heatmap shows the *Igk* locus for pro-B versus B cells. Rows are single cells, columns are 50 kb genomic regions. Blue represents regions where cut fragments are likely coming from H3K27me3, while red represents regions where cut fragments are likely coming from H3K4me1.

We quantify the accuracy of scChIX to select the correct H3K27me3-H3K9me3 cluster pair to mix together. We color each cell by its ground truth label and plot its inferred H3K27me3-H3K9me3 pair (Figure 2b, left). The false discovery rates of scChIX predicting B cells, granulocytes, or NK cells are 10%, 3%, and 1%, respectively (Figure 2b, right). Similarly, scChIX has high specificity and sensitivity.

scChIX assigns each double-incubated cut fragment to either H3K27me3 or H3K9me3 (Figure 2c, Methods). The deconvolved B cell repressive landscapes correspond with their respective ground truth, exemplified in the *Bcl2* locus (Figure 2d) and other loci (Supplemental Figure 4). We also find cell type-specific signal in H3K27me3 (Supplemental Figure 5) and H3K9me3 signal (Supplemental Figure 6).

Our model infers a probability of each cut fragment to be assigned to H3K27me3 or H3K9me3, depending on the genomic location of the cut and the cell (Methods). The inferred probabilities at the *Bcl2* locus reveals the mutual exclusive relationship between H3K27me3 and H3K9me3, where the chromatin state is predominantly H3K9me3, then switches to H3K27me3, and then switches back to H3K9me3 (Figure 2e). For *Bcl2*, these transitions occur at the same location independent of the cell type. However, We find that these transitions can also be cell type-specific, as exemplified by the *Crim1* locus (Figure 2f), where the H3K27me3 region extends further upstream of *Crim1* in NK cells compared with B cells and granulocytes. Our ground truth experiment demonstrates that scChIX can accurately map two histone modifications in single cells, and the inferred probabilities can be biologically interpreted as relationships between the two histone modifications in single cells.

### scChIX reveals joint dynamics of active and repressive chromatin regulation within blood cell types

We next apply scChIX to integrate active (H3K4me1) and repressive (H3K27me3) chromatin states in individual blood cells residing in the bone marrow. We use scChIX to transfer labels and link UMAPs between active and repressive histone modifications (Figure 3a,b) to perform a joint analysis of the two marks.

First, we define cell types from the H3K4me1 sortChIC data by taking the top 150 genes associated with different clusters and use a publicly available scRNA-seq dataset to compare its mRNA abundances of these genes across different blood cell types^19^ (Supplemental Figure 7).

scChIX takes each H3K4me1+H3K27me3 cell and infers the most likely cluster pair (one from H3K4me1, the other from H3K27me3), which systematically transfers cell type labels defined from H3K4me1 onto the H3K27me3 data. (Supplemental Figure 8a). We find that a small minority of double-incubated cells have low-confidence cluster pair predictions. Plotting the cluster pairs onto the H3K4me1+H3K27me3 UMAP confirms that the single-cell assignment produces precise clusters where neighboring cells are likely assigned to the same pair. Low-confidence predictions arise from cells that border between clusters (Supplemental Figure 8b), which we remove from further analysis. Overall, scChIX allows systematic transfer of cell type labels from one histone modification to another.

We next deconvolve the double-incubated cells into their respective histone modification. The UMAPs from H3K4me1 and H3K27me3 show that single-incubated and deconvolved single cells intermingle, suggesting that deconvolution does not produce significant batch effects (Supplemental Figure 9). The deconvolved single cells provide anchors to systematically link one histone modification with another. This joint space reveals a linked landscape between H3K4me1 and H3K27me3, where a location in the UMAP of one histone modification corresponds to a location in the UMAP of another modification. (Figure 3c).

To validate the predicted cell types in both the single- and deconvolved datasets, we compare with data from cell types purified by FACS. For H3K4me1 clusters, we compare with publicly available ChIP-seq^16^. Pearson correlation between ChIP-seq of B cells, erythroids, granulocytes, and NK cells versus sortChIC from single- and double-incubated cells is highest for the predicted cell type (Supplemental Figure 10). Although single-incubated cells have higher correlation with ChIP-seq reference data than deconvolved cells for the matched cell type, the deconvolved cells of the matched cell type consistently had higher correlation with ChIP-seq than unmatched cell types. For H3K27me3 clusters, we use our ground truth sortChIC data purified from FACS. Pearson correlation between sorted sortChIC of B cells, granulocytes, and NK cells versus unsorted bone marrow sortChIC is highest for the predicted cell type (Supplemental Figure 11).

We use the joint landscape to reveal active and repressive histone modification dynamics within cell types. To find differences in chromatin regulation between pro-B cells versus B cells, we select only pro-B or B cells and re-cluster the cells in both H3K4me1 and H3K27me3 separately (Supplemental Figure 12a, b). Using pro-B cell-specific genes, *Pax5*^20^ and *Pten*^21^, we project the H3K4me1 signal at loci overlapping these genes onto both H3K4me1 and H3K27me3 landscapes, confirming a subset of pro-B cells within the B cell population (Supplemental Figure 12c). Similarly, we use marker genes associated with more differentiated B cells, such as *Irf4*^20^, *Igkv3-2* locus^22^, and *Cd72*^23^ to confirm a more differentiated B cell population (Supplemental Figure 12d). Plotting the heatmap of H3K4me1-H3K27me3 assignment probabilities at the *IgK* locus reveals that the chromatin state is repressed in pro-B cells but becomes activated in B cells (Figure 3d), consistent with the progressive activation of the chromatin state during B cell development^22^.

Next, we re-cluster neutrophils to analyze differences in chromatin regulation along pseudotime (Supplemental Figure 13a, Methods). Re-clustering neutrophils in H3K27me3 reveals a shared pseudotime trajectory that varies smoothly between neutrophils in both the H3K27me3 and H3K4me1 landscapes. H3K4me1 levels at the *Retnlg* locus, a marker gene for mature neutrophils^24^ increases along pseudotime, while H3K27me3 levels decreases (Supplemental Figure 13b). The H3K27me3 gene loadings associated with pseudotime consists of a module of *Hox* and other developmental genes (Supplemental Figure 13c-e). Of note, these genes have low levels of mRNA abundances in neutrophils (Supplemental Figure 13f), suggesting that this module is transcriptionally silent. At a locus overlapping the *Hoxa* locus, we find that H3K27me3 was highly marked while H3K4me1 was lowly marked across all neutrophils. Along pseudotime, H3K27me3 further increases, while H3K4me1 further decreases (Supplemental Figure 13c). Our pseudotime analysis suggests that dynamics in histone modifications can occur even in regions associated with lowly expressed genes.

## Discussion

We demonstrate that scChIX can deconvolve multiplexed histone modifications, expanding the number of histone marks that can be profiled in single cells. We generate a ground truth dataset to validate accuracy and precision of scChIX, and apply scChIX to a heterogeneous sample to link active and repressive chromatin regulation at the single-cell level. Our integrated analysis reveals a repressed chromatin signature in immunoglobulin genes in pro-B cells, but become active in B cells. Pseudotime analysis in neutrophils finds that transcriptionally silent regions can have dynamics in repressive histone modification levels. Overall, scChIX unlocks multimodal analysis in antibody-based chromatin profiling and enables joint analysis of distinct histone modifications in single cells.

## Methods

### Animal experiments

Male 13-week-old C57BL/6 mice were used to extract bone marrow cells. Experimental procedures were approved by the Dier Experimenten Commissie of the Royal Netherlands Academy of Arts and Sciences and performed according to the guidelines. Femur and Tibia were extracted, the bones ends were cut away to access the bone marrow which was flushed out using a 22G syringe with HBSS (-ca, -mag, -phenol red, Gibco 14175053) supplemented with Pen-Strep and 1 % FCS. The bone marrow was dissociated and debris was removed by passing it through a 70 μm cell strainer (Corning, 431751). Cells were washed with 25 mm supplemented HBSS before depleting the sample of unnucleated cells using IOTest 3 Lysing solution (Beckman Coulter) following provider instructions. Cells were washed additional 2 times with PBS before processing them by the sortChIC protocol for the histone modifications H3K27me3, H3K4me1 and H3K27me3 + H3K4me1. In case of the fixed cells experiment, isolated cells were resuspended in 70 % ethanol and frozen down at *−*80 °C prior to processing by the sortChIC protocol for the histone modifications H3K27me3, H3K9me3 and H3K27me3 + H3K9me3.

### sortChIC experiments

Cells were processed in 0.5 ml protein low-binding tubes. Following steps are performed on ice. Cells were resuspended in 500 μl Wash Buffer (47.5 ml H2O RNAse free, 1 ml 1 M HEPES pH 7.5 (Invitrogen), 1.5 ml 5M NaCl, 3.6 μl pure spermidine solution (Sigma Aldrich)). Cells were pelleted at 600 g for 3 min and resuspended in 400 μl Wash Buffer 1 (wash buffer with 0.05 % saponin (Sigma Aldrich), protease inhibitor cocktail (Sigma Aldrich), 4 μl 0.5 M EDTA) containing the primary antibody (1:100 dilution for H3K4me1, H3K27me3 and H3K4me1+H3K27me3 Saponin has to be prepared fresh every time as a 10 % solution in PBS). Cells were incubated overnight at 4 °C on a roller, before they were washed once with 500 μl Wash Buffer 2 (wash buffer with 0.05 % saponin, protease inhibitor). Afterwards cells were resuspended in 500 μl Wash Buffer 2 containing Protein A-Micrococcal Nuclease (pA-MNase) (3 ng/ml) and incubated for 1 h at 4 °C on a roller.

Finally, cells were washed an additional 2 times with 500 μl Wash Buffer 2 before passing it through a 70 μm cell strainer (Corning, 431751) and sorting G1 cells based on Hoechst staining on a JAZZ FACS machine into 384 well plates containing 50 nl Wash buffer 3 (Wash buffer containing 0.05 % saponin) and 5 μl sterile filtered mineral oil (Sigma Aldrich) per well. Small volumes were distributed using a Nanodrop II system (Innovadyme).

### Processing of ethanol fixed cells for sortChIC protocol and surface antibody incubation

For the ethanol fixed cells the above described sortChIC protocol was adapted. Wash Buffers were used as described above, except that 0.05 % saponin was exchanged for 0.05 % Tween. Ethanol fixed cells were thawed on ice. Cells were spun at 400 g for 5 min and washed once with 400 μl Wash Buffer 1. Cells were spun again at 400 g and resuspended in 400 μl Wash Buffer 1. Cell suspension was split into 3 samples each having a volume of 400 μl and incubated with one or two antibodies (1:100 dilution for H3K27me3, H3K9me3 and H3K27me+H3K9me3) overnight on a roller at 4 °C. The next day cells were spun at 400 g, washed once with 400 μl Wash Buffer 2 and resuspended in 500 μl Wash Buffer 2 containing pA-MNase (3 ng/ml) and incubated for 1 h on a rotator at 4 °C. Next, cells were spun at 400 g and resuspended in 400 μl Wash Buffer 2 (with addition of 5% blocking rat serum). Surface antibodies were added according to these concentrations and were incubated for 30 min on ice:

| antibody | info | working concentration |
| --- | --- | --- |
| GR1 | A647, anti-mouse Ly-6G/Ly-6C (Gr-1) Antibody, clone: RB6-8C5 | 1:8000 |
| NK1 | A488, anti-mouse NK-1.1 Antibody, clone: PK136 | 1:400 |
| CD19 | BV421, anti-mouse CD19 Antibody, clone: 6D5 | 1:200 |

Finally, samples were washed once with 500 μl Wash Buffer 2 before passing them through a 70 μm cell strainer (Corning, 431751) and sorting on a JAZZ FACS machine, with surface antibody specific gating, into 384 well plates containing 50 nl Wash buffer 3 (Wash buffer containing 0.05 % Tween) and 5 μl sterile filtered mineral oil (Sigma Aldrich) per well. Small volumes were distributed using a Nanodrop II system (Innovadyme).

### Data processing

All DNA libraries were sequenced on a Illumina NextSeq500 with 2×75bp. Standard workflow of sortChIC data process-ing from *singlecellmultiomics* (github.com/BuysDB/SingleCellMultiOmics) was followed^9^ including fastq files demultiplexing, reads mapping with bwa and reads binning into 50kb bins for generating count tables.

### Unmixing scChIX signal

SortChIC generates a vector of cut fragments that map along the genome for each cell. We model the vectors of counts from a double-incubated cell 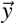 as a mixture of two multinomial distributions, one coming from a cluster *c* of histone modification 1, parametrized by 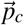, the other from another cluster *d* of histone modification 2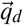 (Supplemental Methods). The log-likelihoood for a mixture of two multinomials is:

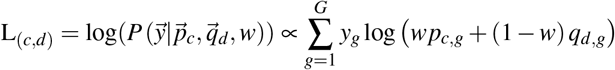

where 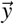 is the number of cuts across the genome for a double-incubated cell, 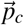 and 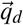 are probabilities for histone modifications 1 and 2 across *G* genomic regions, *w* is the mixing fraction of histone modification 1 in the double-incubated cell. We estimate *w* by maximizing the log-likelihood given 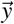, 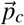, and 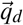

For each double-incubated cell, we infer which pair of multinomials 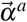 and 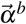 most likely generated the genomic distribution of cuts by calculating the probability that a double-incubated cell comes from a particular pair (*c, d*):

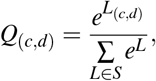

where *L*_*c,d*_ is the likelihood the double-incubated cuts came from mixing cluster *c* from histone modification 1 and cluster *d* from modification 2. *S* is the set of possible pairs of clusters in histone mark 1 (indexed by 1 to C) with clusters in histone mark 2 (indexed by 1 to D):

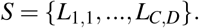

For each double-incubated cell, we select the cluster pair with the highest probability. The mixing fraction *w* is inferred by finding *w* that maximizes *L* for each pair.

We apply Latent Dirichlet Allocation (LDA, Supplemental Methods) to the single-incubated sortChIC data to infer the number of clusters *C* and *D* for histone modifications 1 and 2 as well as the cluster-specific probabilities, 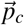 and 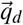.

After assigning each cell to the most likely cluster pair 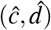, we assign *y*_*i, j*_, the *j*th read mapped to region *g* in cell *i*, to histone mark 1 with probability *P*_*i, j*_:

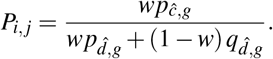

This assignment generates a pair of vectors 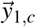 and 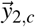 that are linked because they both come from cell *c*. Unmixed counts 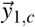 and 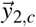 are then be projected back, respectively, onto the space inferred from training data of histone modification 1 and 2. The links between histone modification 1 and 2 are used to transfer labels and create linked UMAPs between the two histone modifications.

## Acknowledgements

We thank Marijn van Loenhout for experimental advice and Reinier van den Linden for FAC-sorting. This work was supported by a European Research Council Advanced grant (ERC-AdG 742225-IntScOmics), Nederlandse Organisatie voor Wetenschappelijk Onderzoek (NWO) TOP award (NWO-CW 714.016.001), the Swiss National Science Foundation, and the Human Frontiers for Science Program. This work is part of the Oncode Institute which is partly financed by the Dutch Cancer Society.

## Author contributions statement

JY, MF, BAdB, and AvO conceived the project. MF developed double-incubation techniques and performed experiments, with help from PZ. JY developed and applied statistical methods, with help from MF and BAdB. BAdB wrote the sortChIC preprocessing pipeline, with help from MF and JY. JY, MF, and AvO analyzed the data. JY, MF, AvO wrote the manuscript, with input from PZ and BAdB.

