## Supplemental_Information for "Deconvolving multiplexed histone modifications in single cells"

**This PDF includes:**

- Supplemental methods
- Supplemental figures and legends 1-13

### 1 Statistical framework for unmixing scChIX data

#### 1.1 Overview of the statistical model

Our goal is to unmix sortChIC data from cells incubated with two different antibodies into its separate histone modifications. SortChIC generates a vector of cut fragments that map along the genome for each cell. For each cell, we assume the distribution of discrete cut fragments in the genome comes from a multinomial distribution. Each genomic region $g$ has a probability $f_g$ to generate a cut, where $\sum_g f_g = 1$. Each cell generates a total of $\sum_g y_g = N$ cuts.

$$\vec{y} \sim \text{Multinomial}(\vec{f}, N)$$

In a double immunocleavage experiment, we assume that $\vec{f}$ come from a mixture of two multinomials $\vec{p}$ and $\vec{q}$ (one from each histone modification). The two multinomials are mixed with a mixing fraction $w$ such that for a region $g$:

$$f_g = w p_g + (1-w) q_k$$

The multinomial log-likelihood of a cell with cuts across the genome $\vec{y}$ mixed from multinomials is therefore:

$$\mathsf{L} = \log(P(\vec{y}|\vec{p},\vec{q},w)) \propto \sum_{g=1}^{G} y_g \log(wp_g + (1-w)q_g),$$

where genomic regions are defined by 50 kb bins in the genome.

Since the sortChIC experiments are done in complex samples with different cell types, $\vec{p}$ and $\vec{q}$ are cell type specific. We use the single-incubated data as training to infer the number of clusters for each histone modification ($|C|$ and $|D|$), respectively) as well as the cell type-specific probabilities $\vec{p}_c$ and $\vec{q}_d$. The probability that a duo-cut cell comes from a particular pair $(c,d)$:

$$Q_{(c,d)} = \frac{e^{L_{(c,d)}}}{\sum\limits_{L \in S} e^{L}},$$

where $L_{c,d}$ is the likelihood the double-incubated cuts came from mixing cluster $c$ from histone modification 1 and cluster $d$ from modification 2. $S$ is the set of possible pairs of clusters in histone mark 1 (indexed by 1 to C) with clusters in histone mark 2 (indexed by 1 to D):

$$S = \{L_{1,1}, ..., L_{C,D}\}.$$

For each pair, we use the Brent method implemented in R (*optim*) to find $w$ that minimizes the negative log-likelihood. We then select the pair $(c,d)$ with the highest probability $Q$.

#### 1.2 Generating the possible pairs of genomic profiles

We use Latent Dirichlet Allocation (LDA), a matrix decomposition model based on a hierarchy of multinomials, to model the relative frequencies of cuts in each genomic region in single cells. LDA can be thought of as a discrete analogue to principal component analysis.

To generate the genomic location of the $j$th read for cell $i$:

Choose a topic $z_{i,j}$ by sampling from the cell-specific distribution of topics:

$$\vec{U}_i \sim \text{Dirichlet}(\alpha)$$

$$z_{i,j} \sim \text{Multinomial}(\vec{w}_i, 1)$$

Choose genomic region $w_{i,j}$ by sampling from the topic-specific distribution of genomic regions:

$$\vec{V}_k \sim \text{Dirichlet}(\delta)$$

$$w_{i,j} \sim \text{Multinomial}(\vec{V}_{z_{i,j}}, 1)$$

The Dirichlet distributions are priors to prevent overfitting when there are few cuts in the region. We used the LDA model implemented by the topicmodels R package, using the Gibbs sampling implementation with hyperparameters $\alpha = 1.67$, $\delta = 0.1$, where K is the number of topics[25].

We cluster cells into cell types by analyzing distribution of topic weights across cells, which has a bimodal distribution to indicate cells that belong to one cluster versus not refsection:DefineClusters. We estimate $\vec{p}_c$ and $\vec{q}_d$ for each cluster in histone modification 1 $\{\vec{p}_1, \vec{p}_2, ..., \vec{p}_C\}$ and modification 2 $\{\vec{q}_1, \vec{q}_2, ..., \vec{q}_D\}$ by averaging the estimated probabilities across cells assigned to each cluster:

$$\vec{p}_c = \frac{1}{|A|} \sum_{i=1}^{|A|} \sum_{k=1}^{K} V_{g,k} U_{k,i}$$

where $A$ is the set of cells that belong to cluster $c$.

#### 1.3 Probabilistic assignment of double-incubated reads to their respective histone modification

After assigning each cell to the most likely cluster pair $(\hat{c}, \hat{d})$, we assign $y_{i,j}$, the $j$th read mapped to region $g$ in cell $i$, to histone mark 1 with probability $P_{i,j}$:

$$P_{i,j} = \frac{w p_{\hat{c},g}}{w p_{\hat{d},g} + (1-w)\, q_{\hat{d},g}}.$$

Reads are therefore assigned to the other histone mark 2 with probability $1 - P_{i,j}$

The probabilities $\vec{P}$ are used to split a BAM file of a double-incubated cell into two BAM files, one for each histone modification. When splitting reads at the BAM level, some genomic regions may not have been considered in the model (e.g., regions with few reads in the training data). Probabilities for reads in a missing region $P'_x$ at position $x$ are imputed by linear interpolation between the nearest left $P_{x_l}$ at $x_l$ and right genomic bin $P_{x_r}$ at $x_r$:

$$P'_x = P_{x_l} + (x - x_l) \frac{P_{x_r} - P_{x_l}}{x_r - x_l}$$

### 2 Inferring topic weights in the LDA model

To infer topic weights in the LDA framework, we use a collapsed Gibbs sampling algorithm implemented in the `topicmodels` R package[25]. Briefly, the posterior distribution of the LDA model is inferred from the data by sampling from the update equation:

$$p(z_{d,n} = k | \vec{z}_{-d,n}, \vec{w}, \alpha, \delta) \propto \frac{n_{d,k} + \alpha_k}{\sum_i^K n_{d,i} + \alpha_i} \frac{v_{k,w_{d,n}} + \delta_{w_{d,n}}}{\sum_{j \in \{J\}} v_{k,j} + \delta_j} \quad \text{(Supplemental Equation 1)}$$

Where:

- $z_{d,n}$ is the topic assignment for cell $d$ used to generate cut $n$. $\vec{z}_{-d,n}$ denotes $z_{d,n}$ with the assignment $z_{d,n}$ removed (i.e., the $n$th cut from cell $d$ that will have its topic assignment updated is removed),
- $n_{d,k}$ is the number of times cell $d$ uses topic $k$ across all genomic regions,
- $v_{k,w_{d,n}}$ is the number of times topic $k$ uses genomic region $w_{d,n}$ across all cells,
- $\vec{w}$ is the vector of cuts in the genome across all cells,
- $\alpha$ is the Dirichlet parameter, which can be interpreted as a pseudocount for cell to topic counts,
- $\delta$ is the Dirichlet parameter, which can be interpreted as a pseudocount for topic to genomic region counts.

### 3 Defining cell clusters using LDA topic weights

LDA decomposes the count matrix into two matrices: a cell to topic and a topic to genomic region matrix. The cell to topic matrix assigns each cell to a vector of topic weights indicating a grade of membership of a cell across topics. For each cell type-specific topic, plotting the distribution of topic weights across cells often reveals a bimodal distribution, suggesting a natural cutoff for assigning a cell to a cluster. We fit a Gaussian mixture model to each distribution to estimate the threshold at which the value of the two fitted Gaussian disributions are equal. We assign cells to clusters if the topic weights for the cell is greater than the threshold $t$.

### 4 Transferring cell type labels derived from one histone modification to another modification

To transfer labels from one sortChIC data targeting histone modification 1 to modification 2, we unmix the double-incubated cuts $\vec{y}_i$ for each cell $i$ into $\vec{y}_{i,1}$ and $\vec{y}_{i,2}$. We project respectively $\vec{y}_{i,1}$ and $\vec{y}_{i,2}$ onto the latent space of histone modification 1 and 2 inferred from LDA. We assign cell $i$ to cell type $c$ if the topic corresponding to cell type $c$ has a weight $\vec{U}_{1,i} > t$. This threshold $t$ comes from fitting a Gaussian mixture model to define which cells in the training data belong to cluster $c$ (see Section 3). Since cell $i$ has a link in both histone modification 1 and 2, the label from $i$ inferred from histone modification 1 is then transferred to histone modification 2. Single-incubated cells neighboring cell $i$ in histone modification 2, quantified by thresholding cell-to-topic weights in $U_2$, are inferred to have the same label as cell $i$.

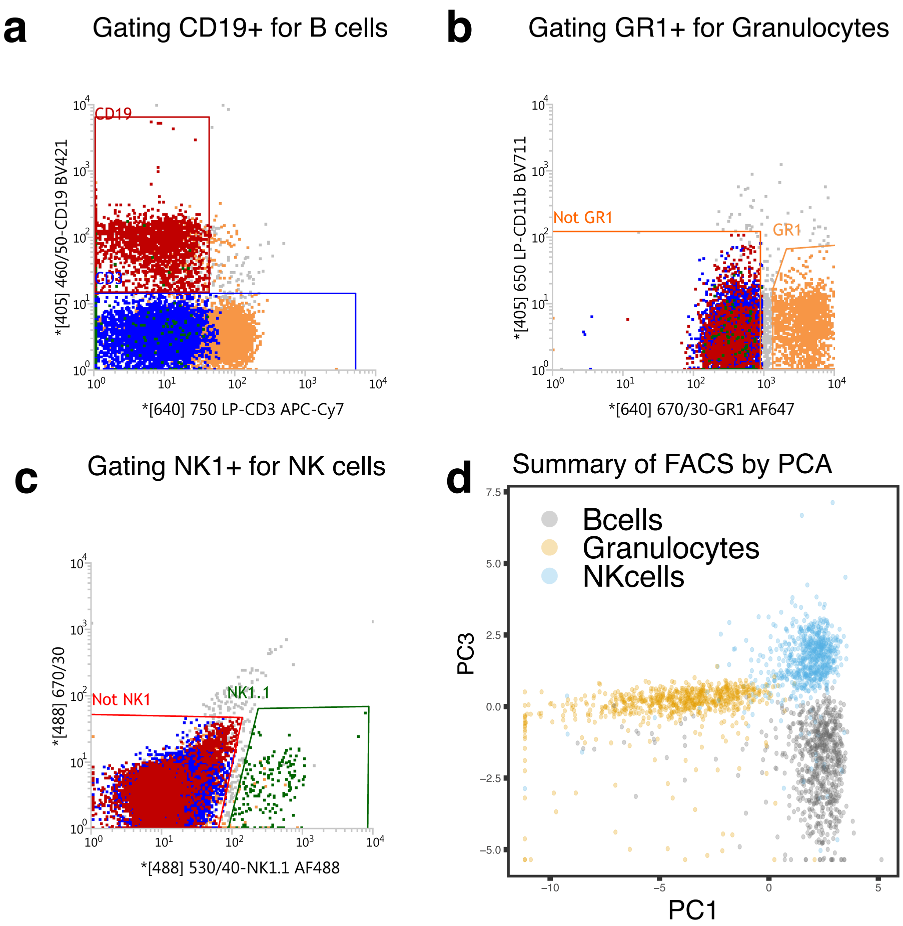


**Supplemental Figure 1.** **FACS gating strategies of fixed cells with surface antibodies. a** Gating strategy of CD19$^{+}$ cells for B cells sorting. **b** Gating strategy of GR1$^{+}$ cells for Granulocytes sorting. **c** Gating strategy of NK1$^{+}$ cells for NK cells sorting. **d** PCA representation of FACS parameters for the three sorted cell types.

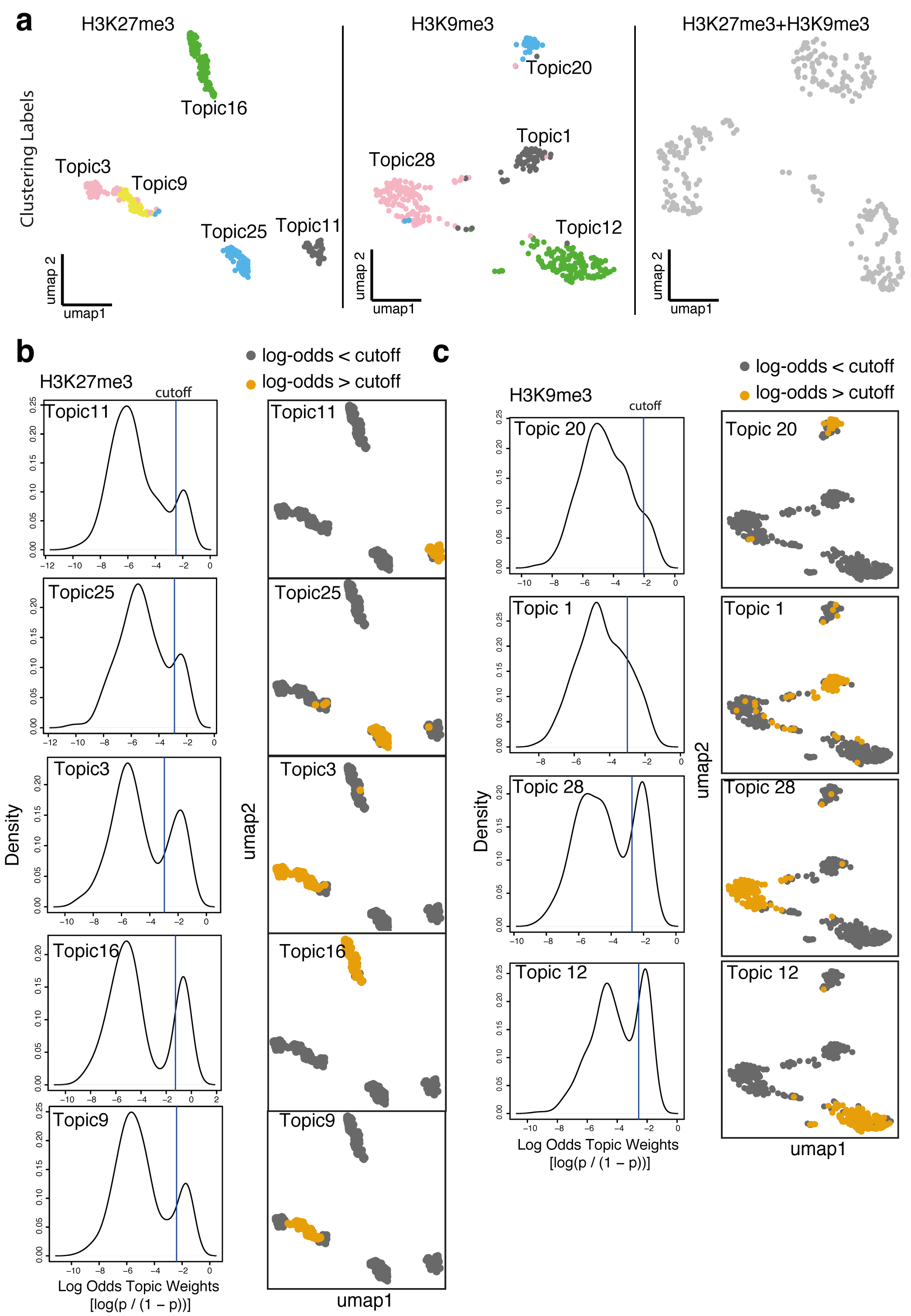


**Supplemental Figure 2.** **LDA infers cell type-specific clusters in sortChIC data.** **a** UMAP of H3K27me3 (left), H3K9me3 (middle), and H3K27me3+H3K9me3 (right) single-cell histone modification landscapes. For the single-incubated datasets, the clusters are colored and labeled as topics. Double-incubated cells are in grey because cell type labels will be inferred from the single-incubated datasets. **b** Density plot (left) of topic weights for different cell type-specific topics across cells in H3K27me3. Vertical line shows the cutoff above which cells are clustered into the topic. UMAP visualization of cells (right) colored by whether topic weights exceed the cutoff. **c** same to (b) but showing clusters in the H3K9me3 signal.

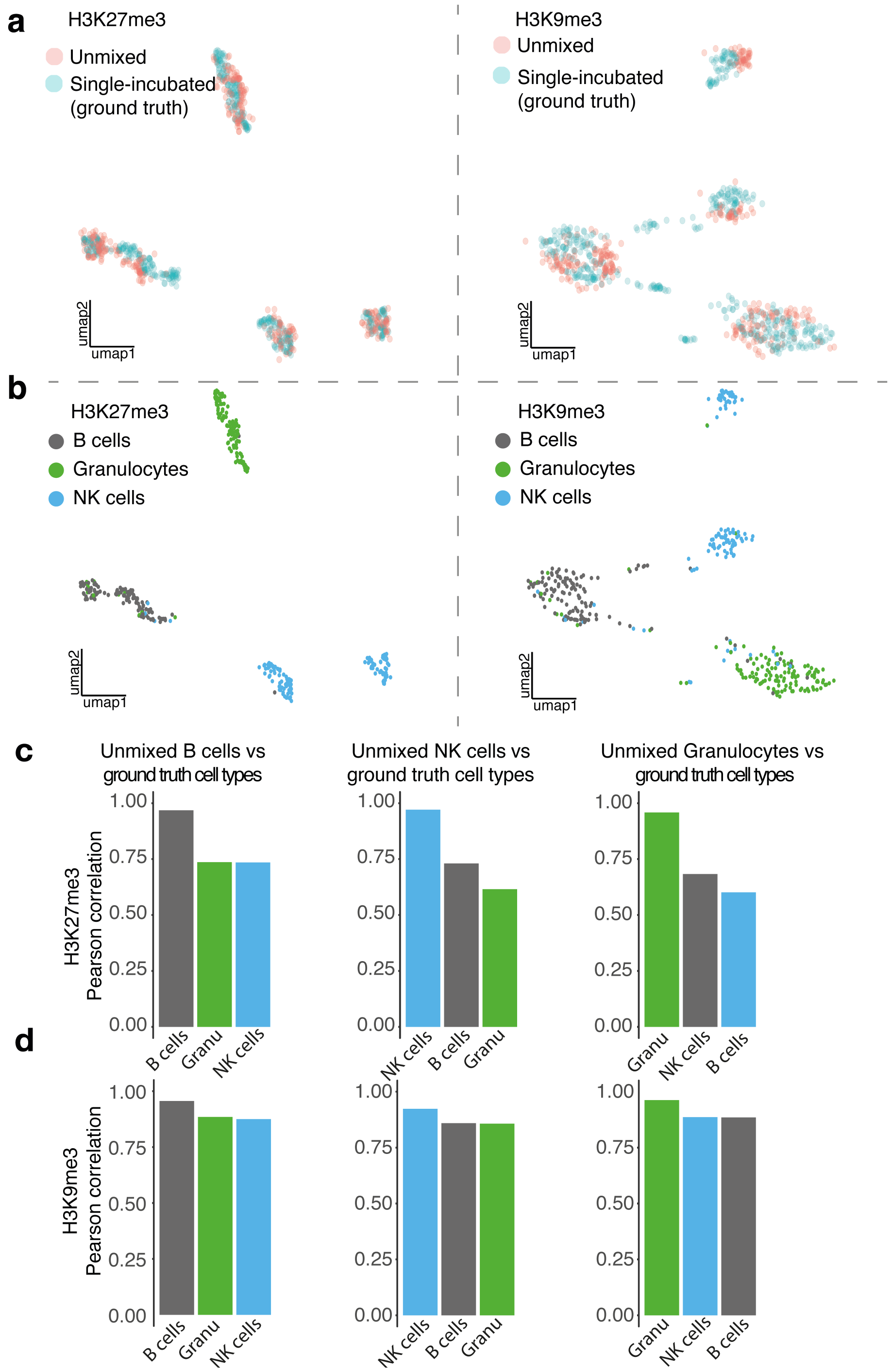


**Supplemental Figure 3. scChIX accurately deconvolves double-incubated signal into their respective histone modifications. a, b** UMAP representation of H3K27me3 (left) and H3K9me3 (right) data colored by unmixed or single-incubated cells (a) or ground truth cell type labels defined by FACS (b). **c, d** Genome-wide Pearson correlation between H3K27me3 (c) and H3K9me3 (d) H3K27me3+H3K9me3 signal versus ground truth sortChIC purified by FACS.

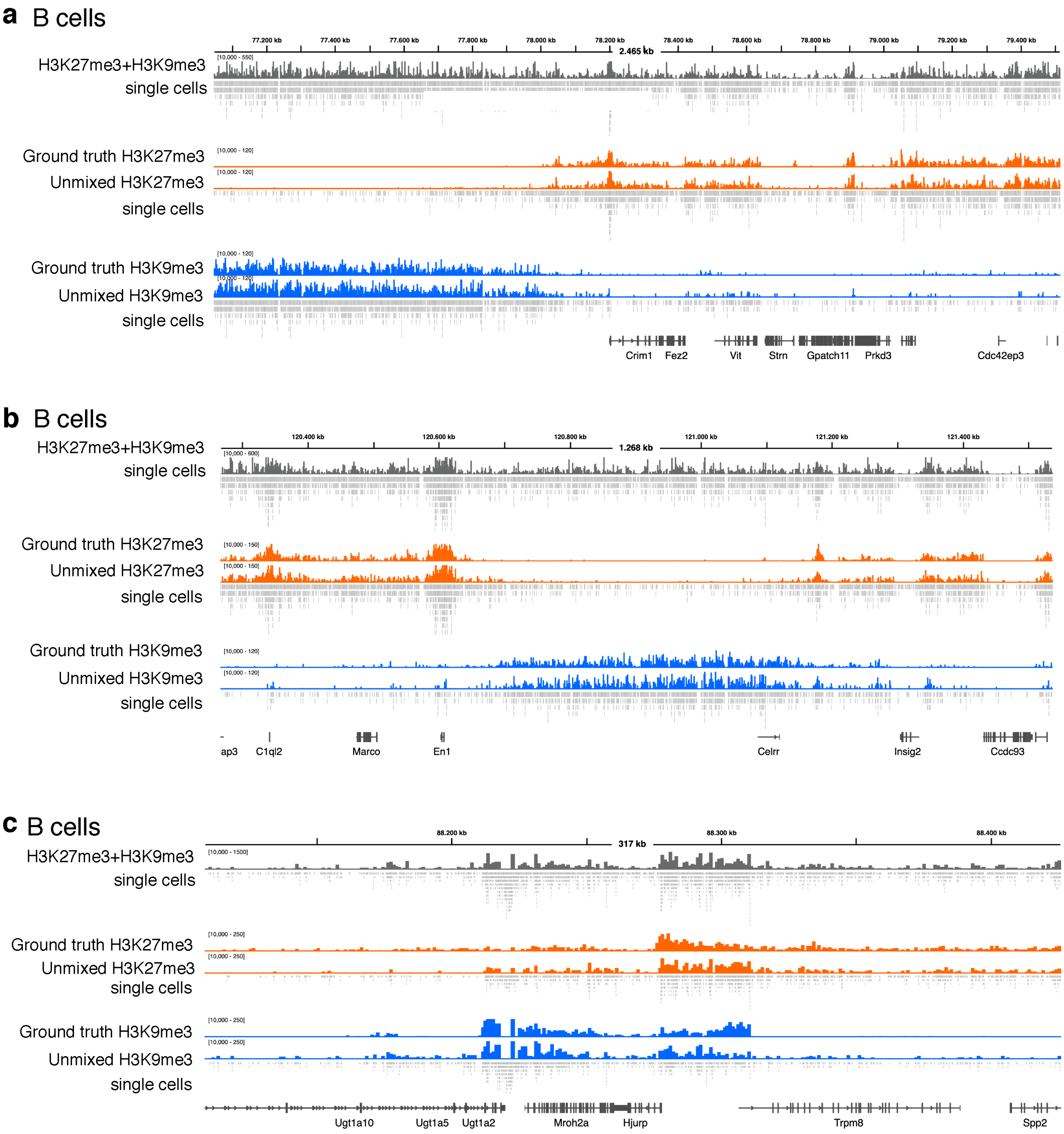


**Supplemental Figure 4. scChIX deconvolves signal that correspond to the ground truth. a-c** Coverage tracks for B cells visualizing the H3K27me3+H3K9me3, deconvolved H3K27me3 or H3K9me3, and ground truth H3K27me3 or H3K9me3 histone modification levels for three different genomic regions. Double-incubated signal in grey, H3K27me3 single, and unmixed signal in orange, and H3K9me3 single and unmixed signal in blue. Under each coverage track are cut fragments of single cells.

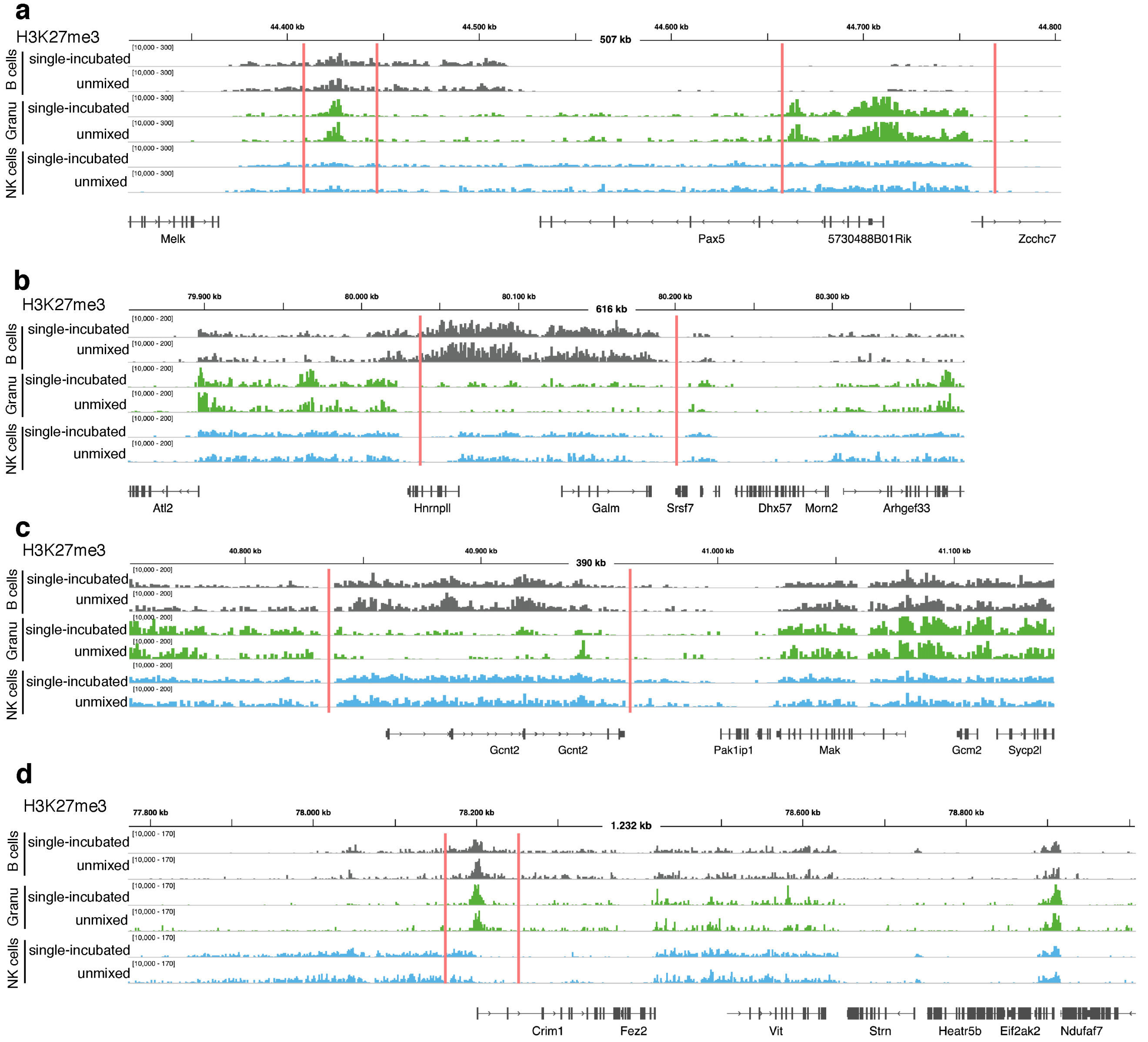


**Supplemental Figure 5. scChIX deconvolves signal at cell type-specific regions for H3K27me3 mark. a** H3K27me3 coverage tracks showing the region around *Pax5* for the ground truth H3K27me3 pseudobulk signal from single-incubated cells and for the deconvolved H3K27me3 pseudobulk signal from double-incubated cells for three cell types: B cells (grey), granulocytes (green), and NK cells (blue). **b** Same as in (a), but for regions around *Hnrnpll* and *Galm*. **c** Same as in (a) but for a region around *Gcnt2*. **d** Same as in (a) but for a region around *Crim1*

.

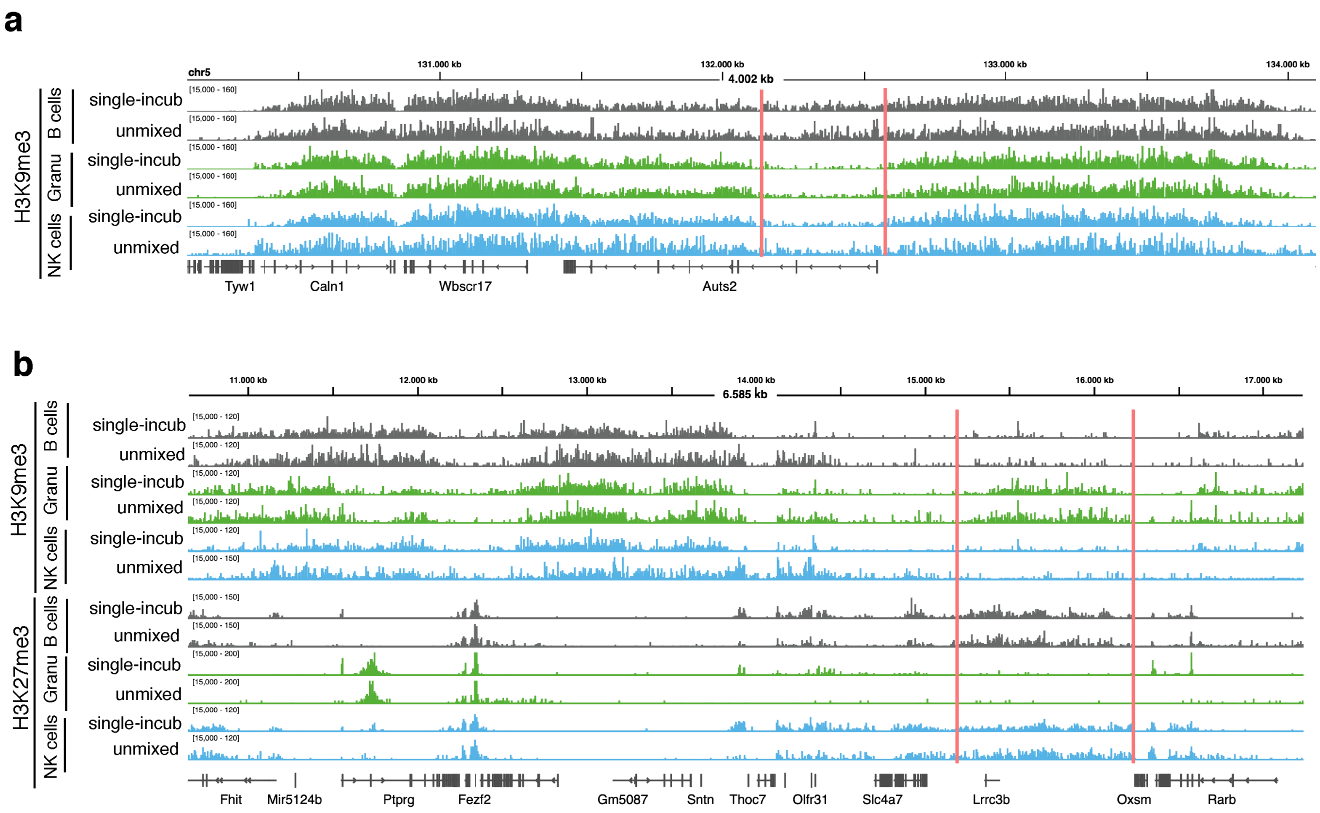


**Supplemental Figure 6. scChIX deconvolves signal at cell type-specific regions for H3K9me3 mark. a** H3K9me3 coverage tracks showing the region around *Auts2* for ground truth H3K9me3 (single-incubated) and for the unmixed H3K9me3 (unmixed) for B cells (grey), granulocytes (green) and NK cells (blue), respectively. **b** Same as (a), but for a region around *Lrrc3b*.

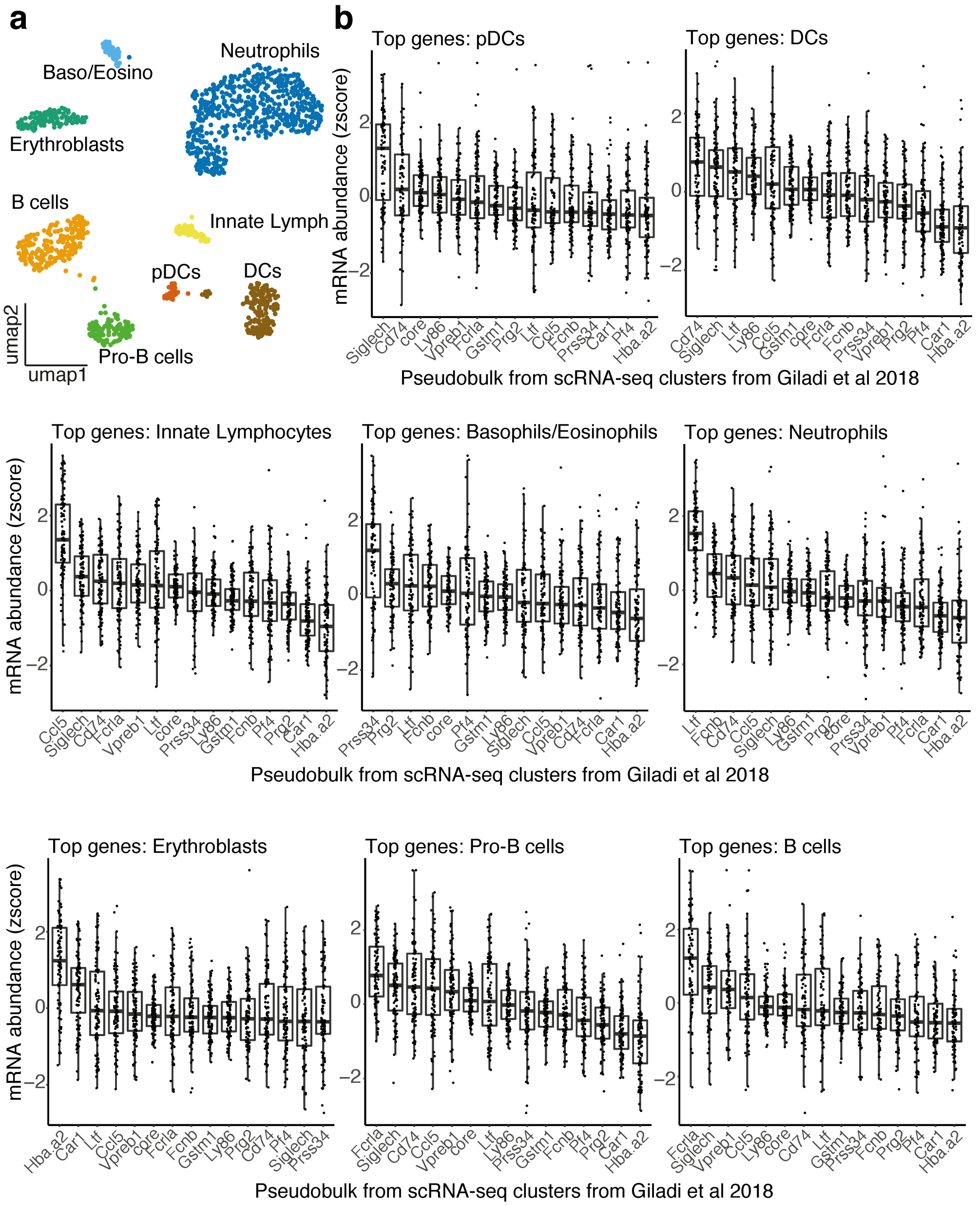


**Supplemental Figure 7. Cell typing clusters from H3K4me1 in mouse bone marrow. a** UMAP of H3K4me1 sortChIC data, cells colored by cell type. **b** Boxplots of mRNA abundances of genes associated with the top 150 cluster type-specific regions (eight boxplots, each for a cluster defined by H3K4me1). mRNA abundances come from pseudobulks of scRNA-seq[19]. Name of pseudobulks is associated with marker genes that define the cell type: *Ltf*=neutrophils, *Siglech*=plasmacytoid dendritic cells, *Cd74*=conventional dendritic cells, *Ccl5*=innate lymphocytes, *Hba-a2*=erythroblasts, *Fcrla*=B cells, *Prss34*=Basophils, *Prg2*=Eosinophils.

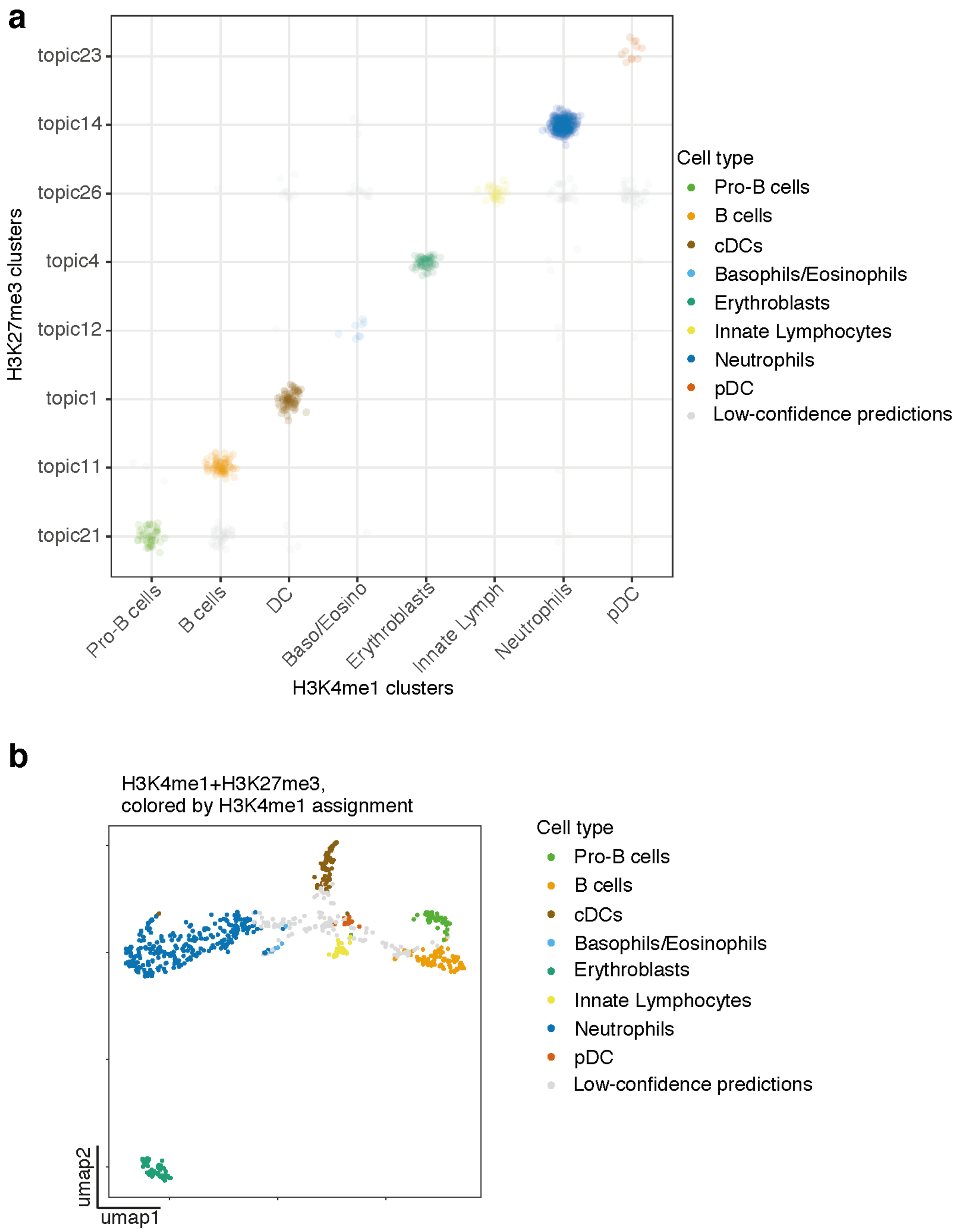


**Supplemental Figure 8. Inferring cluster pairs from H3K4me1+H3K27me3 transfers cell type labels. a** Assignment plot showing individual H3K4me1+H3K27me3 cells (represented as dots) assigned to a pair of topics (x-axis labels are H3K4me1 clusters, named by their associated cell type, while y-axis are H3K27me3 clusters). Cells along the diagonal are high-confidence predictions that match a H3K4me1 cluster with a H3K27me3 topics, and are colored by the H3K4me1-derived cell type labels. **b** UMAP of H3K4me1+H3K27me3 sortChIC. Cells are colored by their cell type inferred from cluster pairs. Low-confidence predictions are colored in grey.

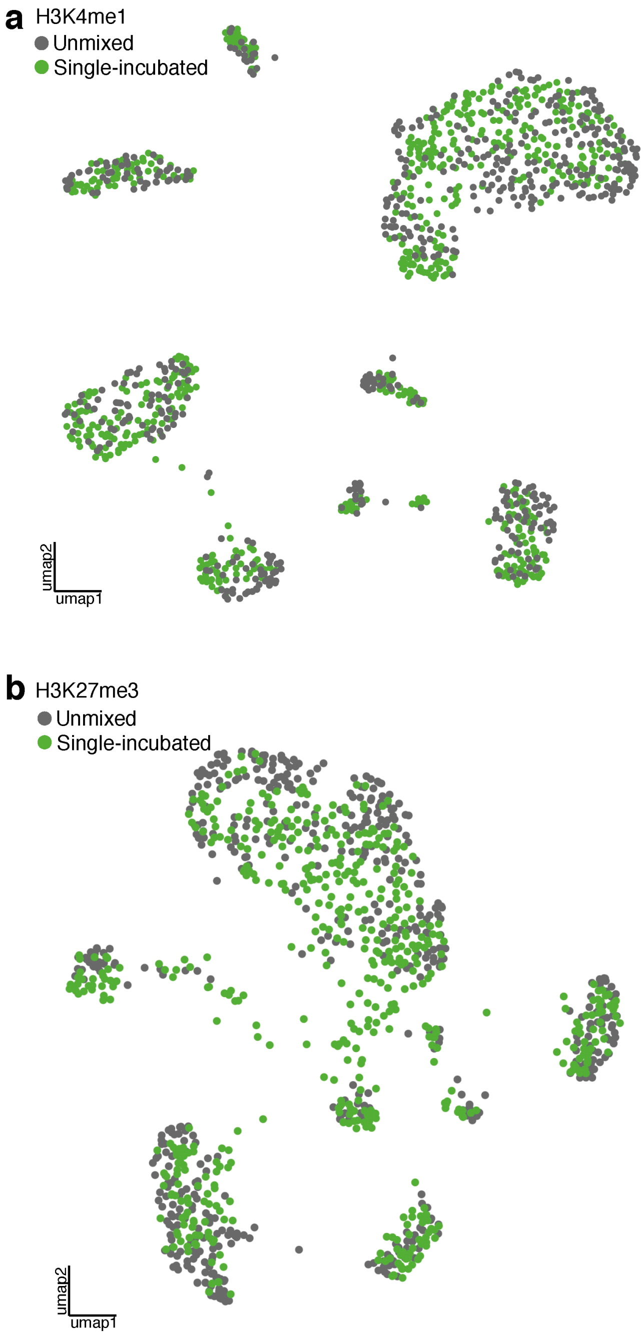


**Supplemental Figure 9. Deconvolved H3K4me1+H3K27me3 cells intermingle with single-incubated cells. a,b** UMAP representation of H3K4me1 (a) and H3K27me3 (b). Cells are colored by whether the epigenome was generated by single-incubation or by unmixing by scChIX.

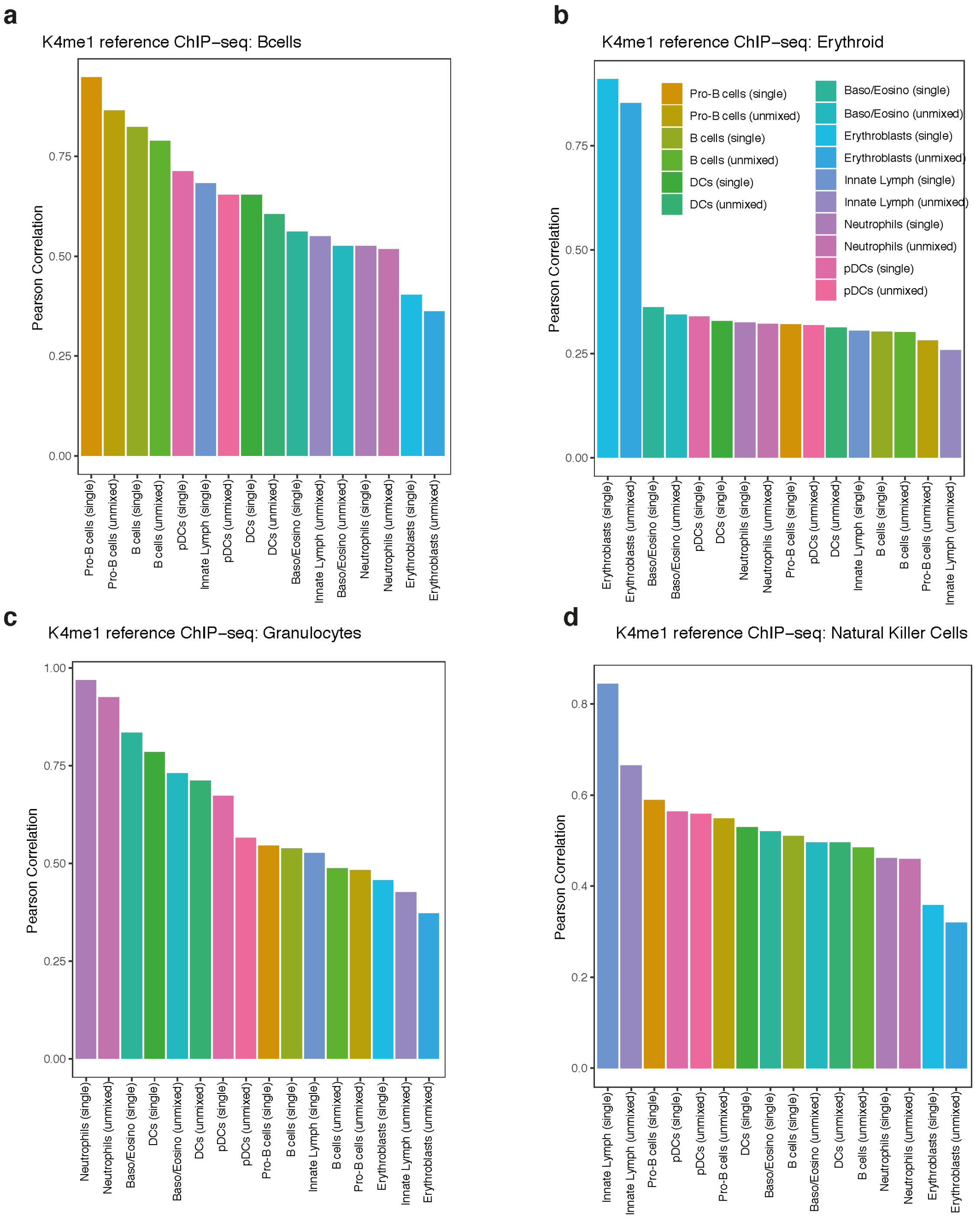


**Supplemental Figure 10. Histone modification signal of deconvolved cell types correlates with publicly available H3K4me1 ChIP-seq data. a** Pearson correlation between publicly available[16] B cell H3K4me1 ChIP-seq versus different H3K4me1 sortChIC derived pseudobulks. Single: epigenomes generated by single incubation, unmixed: epigenomes deconvolved by scChIX. **b** Same as (a) but between erythroid H3K4me1 ChIP-seq versus sortChIC. **c** Same as (a) but between granulocyte H3K4me1 ChIP-seq versus sortChIC. **d** Same as (a) but between natural killer cell H3K4me1 ChIP-seq versus sortChIC.

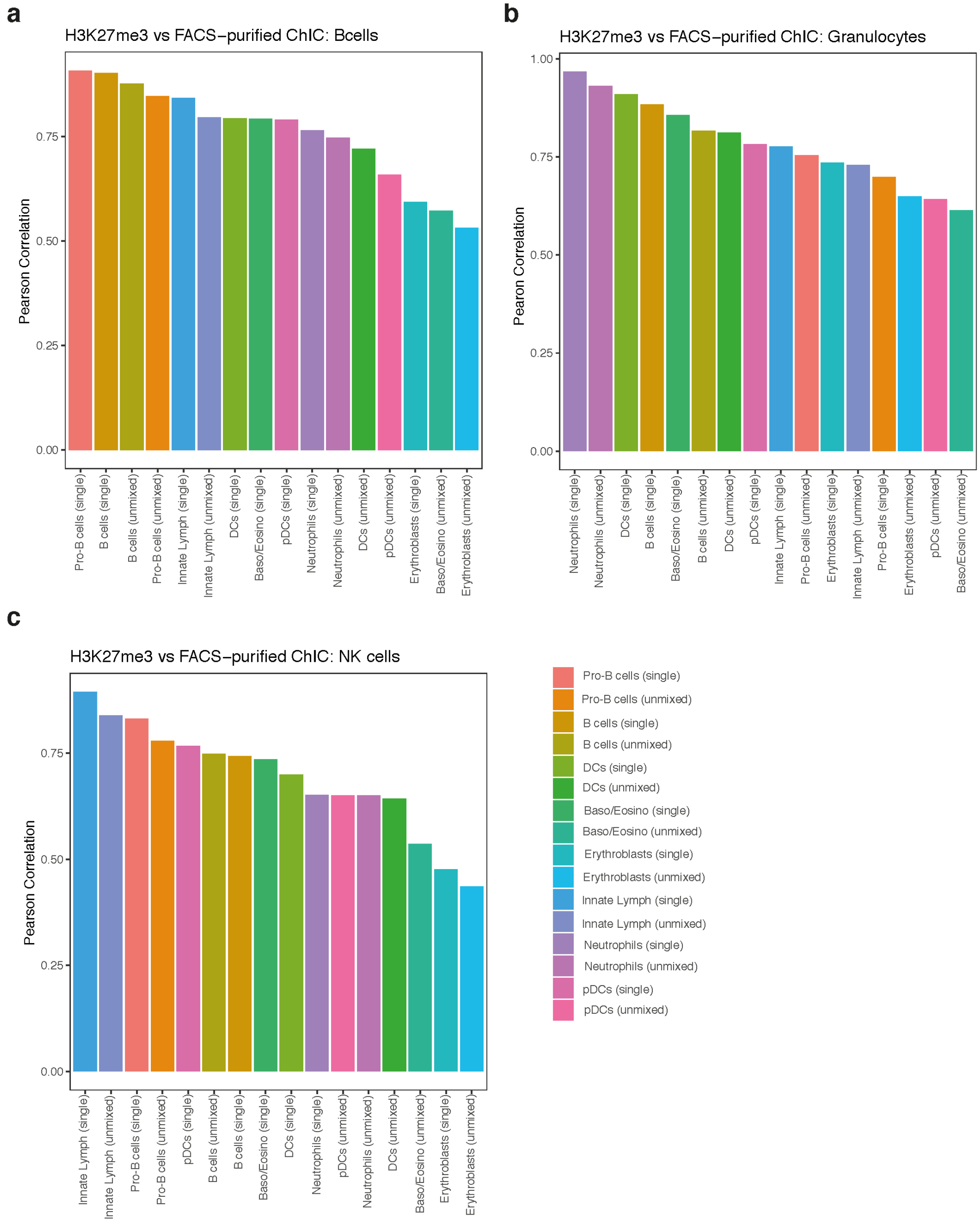


**Supplemental Figure 11. Histone modification signal of deconvolved cell types correlations with H3K27me3 sortChIC of FACS-purified cell types. a** Pearson correlation between H3K27me3 sortChIC from FACS-purified B cells (i.e. pseudobulk from our ground truth dataset) and H3K27me3 sortChIC derived from pseudobulks of whole bone marrow. Single: epigenomes generated by single incubation, unmixed: epigenomes deconvolved by scChIX. **b** Same as (a) but between FACS-purified granulocytes H3K27me3 sortChIC (pseudobulk) versus H3K27me3 sortChIC pseudobulks. **c** Same as (a) but between natural killer cell H3K27me3 sortChIC (pseudobulk) versus H3K27me3 sortChIC pseudobulks.

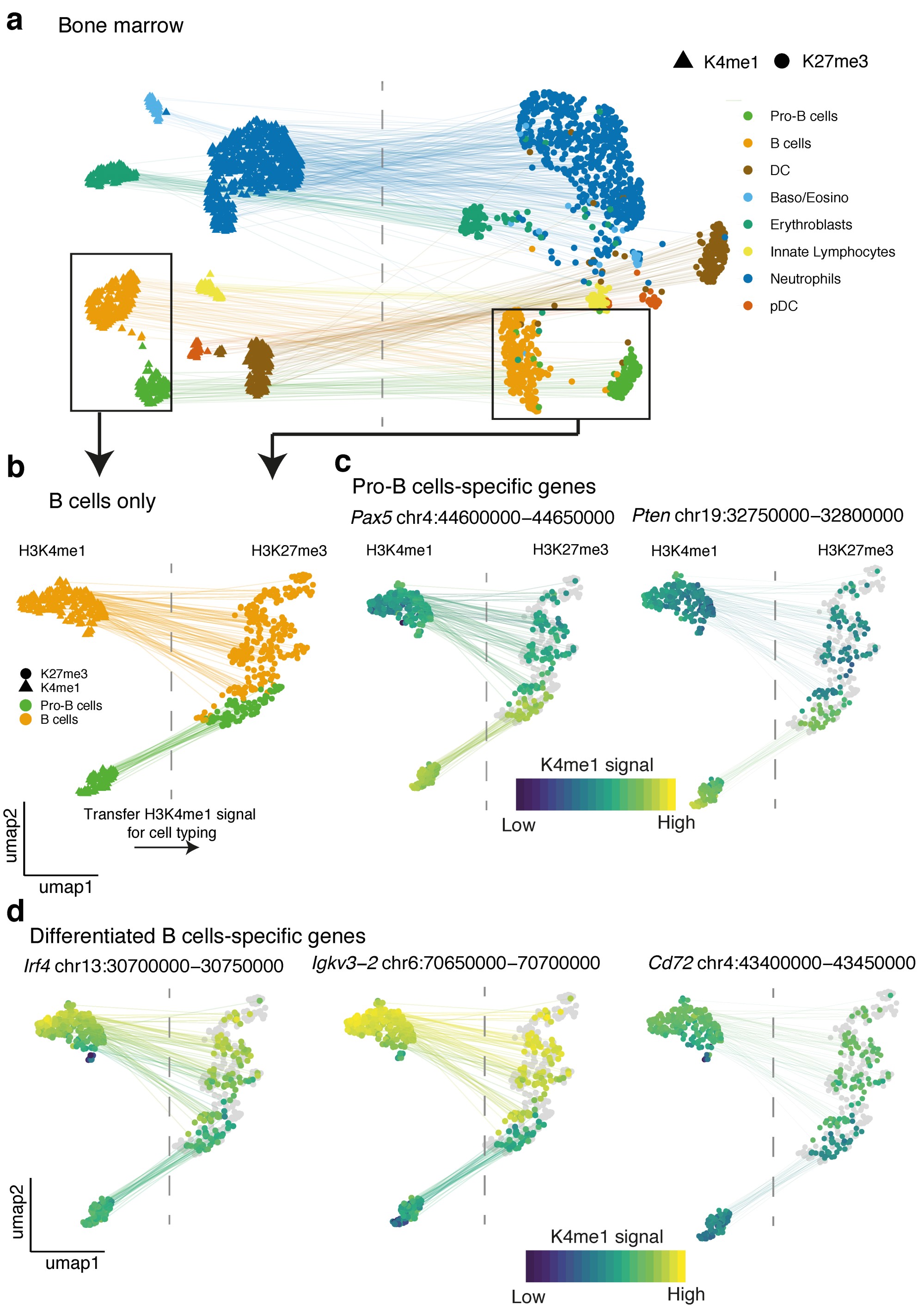


**Supplemental Figure 12. Re-clustering on B cells reveals heterogeneity within B cells. a** UMAP visualization of H3K4me1 and H3K27me3 (single signal and unmixed signal), colored by cell types derived from H3K4me1 and transferred to H3K27me3. Black rectangle indicates the B cell population used to re-cluster in (b,c,d). **b** UMAP of pro-B and B cells only. **c,d** Projection of H3K4me1 signal of marker genes for pro-B (c) or for differentiated B cells (d). H3K4me1 signal is measured in all cells of the H3K4me1 UMAP (i.e. both single- and double-incubated have H3K4me1 signal in the H3K4me1 UMAP). Double- (colored) but not single-incubated (grey) cells have H3K4me1 signal in the H3K27me3 UMAP.

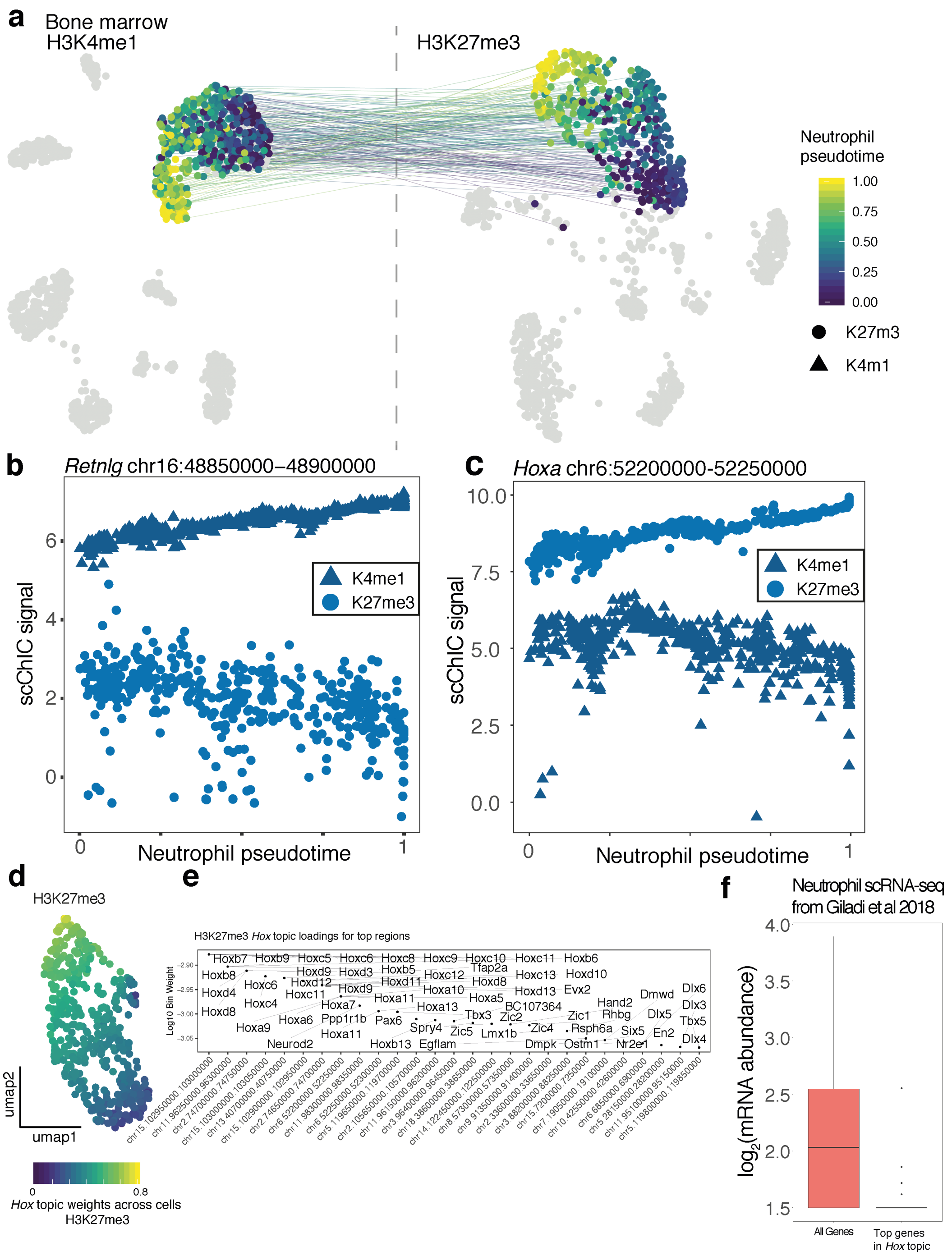


**Supplemental Figure 13.** **H3K4me1 and H3K27me3 signal of during neutrophil maturation.** **a** UMAP visualization of H3K4me1 and H3K27me3, lines join H3K4me1 and H3K27me3 UMAPs of double-incubated neutrophils. Heterogeneity within neutrophils are colored as neutrophil pseudotime. **b** H3K4me1 and H3K27me3 modification levels at the *Retnlg* (a mature neutrophil marker gene) locus along neutrophil pseudotime. **c** H3K4me1 and H3K27me3 modification levels at the *Hoxa* along neutrophil pseudotime. **d** UMAP of H3K27me3 signal across single cells colored by weights of a topic containing high H3K27me3 levels at many *Hox* and developmental gene loci (*Hox* topic). **e** Topic weights of the top 150 genes associated with loci in the *Hox* topic for H3K27me3. **f** Neutrophil mRNA abundance of genes in the *Hox* topic compared to other genes derived from publicly available scRNA-seq data[19].
